# Assembly Mechanism of Mucin and von Willebrand Factor Polymers

**DOI:** 10.1101/2020.03.08.982447

**Authors:** Gabriel Javitt, Lev Khmelnitsky, Lis Albert, Nadav Elad, Tal Ilani, Ron Diskin, Deborah Fass

## Abstract

The respiratory and intestinal tracts are exposed to physical and biological hazards accompanying the intake of air and food. Likewise, the vasculature is threatened by inflammation and trauma. Mucin glycoproteins and the related von Willebrand factor (VWF) guard the vulnerable cell layers in these diverse systems. Colon mucins additionally house and feed the gut microbiome. Here we present an integrated structural analysis of multimerized intestinal mucin MUC2. Our findings reveal the shared mechanism by which complex macromolecules responsible for blood clotting, mucociliary clearance, and the intestinal mucosal barrier form protective polymers and hydrogels. Specifically, cryo-electron microscopy and crystal structures show how disulfide-rich bridges and pH-tunable interfaces control successive assembly steps in the endoplasmic reticulum and Golgi. Remarkably, a densely O-glycosylated mucin domain performs a specific organizational role in MUC2. The mucin assembly mechanism and its adaptation for hemostasis provide the foundation for rational manipulation of barrier function and coagulation.

## INTRODUCTION

In the respiratory tract, secreted mucin glycoproteins are rolled along the airway surface by the movement of cilia to flush out entrapped pathogens and particulate matter (Bustamante-Marin et al., 2017). In the intestines, mucins support the gut microbiome (Johansson et al., 2011) while keeping intestinal contents at a safe distance from the epithelium and lubricating the mucosal surface (Figure 1A). To carry out these essential functions, gel-forming mucins are cross-linked into extended disulfide-linked architectures. The respiratory mucins form bundles of linear covalent polymers (Thornton et al., 2018), and it was proposed that the intestinal mucins form a stacked hexagonal network (Ambort et al., 2012; Nilsson et al., 2014). The non-mucin molecule VWF, produced by endothelial cells and platelets to function in hemostasis and thrombosis (Springer, 2014), evolved by exploiting aspects of the mucin disulfide-mediated assembly mechanism despite its distinct physiological context (Figure 1A).

**Figure 1.**
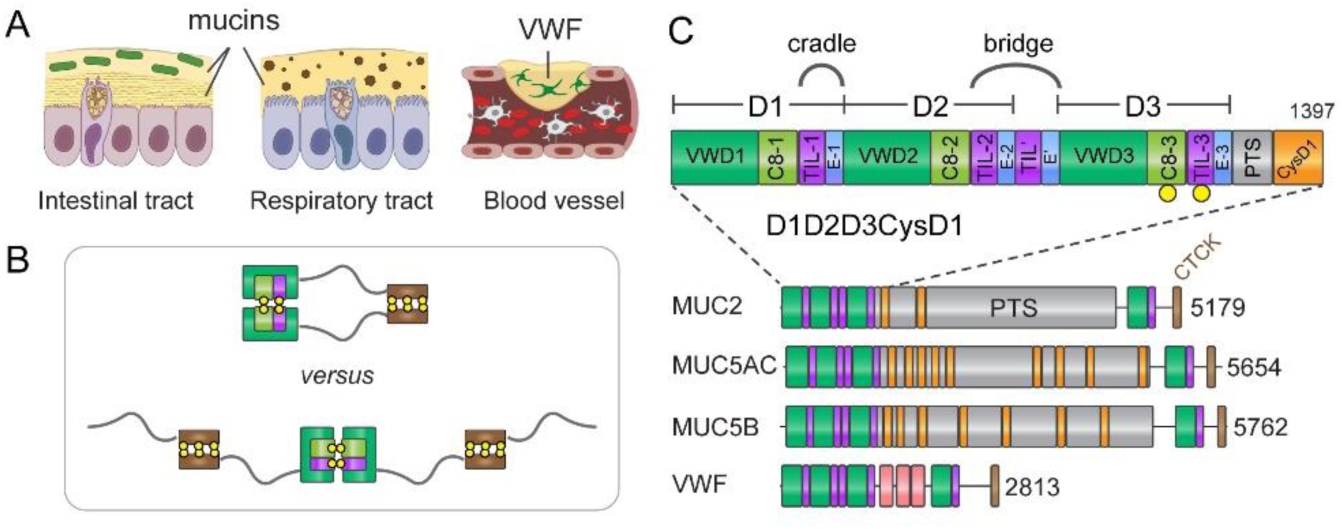
Functions and primary structures of mucins and von Willebrand factor. (A) Functions of mucins and VWF. Green rods: bacteria of the intestinal microbiome; brown hexagons: particulate matter in the respiratory tract; green cells associated with VWF: activated platelets. (B) Flexible molecules that associate by their termini could form either defined oligomerization states (top) or polymers (bottom). Relevant domains are displayed and color-coded as in panel C. (C) Domain organization of gel-forming mucins and VWF. Protein lengths in amino acids are shown to the right. The MUC2 D1D2D3CysD1 region is expanded to show all domains (Nilsson et al., 2014). Positions of cysteines forming intersubunit disulfides during mucin assembly are indicated by yellow balls. The CTCK domain mediates carboxy-terminal disulfide-linked dimerization (Zhou and Springer, 2014). Human MUC2 has a PTS stretch of more than 2300 amino acids with 60% threonine and 20% proline content.

Mucin and VWF assembly begins with formation of disulfide-mediated dimers linked by their carboxy termini in the endoplasmic reticulum (ER), followed by higher-order covalent multimerization *via* their amino termini later in the secretory pathway (Perez-Vilar et al., 1996; Perez-Vilar et al., 1998; Wagner et al., 1984). An unresolved aspect of multimer assembly is how molecules with a tendency to associate at each end form polymers or networks rather than closed dimers (Figure 1B). To date, no explanation for ensuring polymerization has been presented, though the problem has long been appreciated in the context of VWF assembly (Huang et al., 2008; Zhou et al., 2011). Furthermore, it is not obvious, considering their extensive homology (Figure 1C), how the respiratory and intestinal mucins could form different multimeric structures (linear polymers *vs*. hexagonal networks). The current study investigates the mechanism and product of intestinal mucin assembly and relates this mechanism to respiratory mucins and VWF. Illuminating mucin assembly is important because it provides the basis for addressing mucin-related diseases such as cystic fibrosis and colitis (Hansson, 2019), for manipulating the environment of the gut microbiome (Gouyer et al., 2015; Schroeder, 2019), for augmenting the primary host barrier against infections (Linden et al., 2008; Desai et al., 2016), and for enhancing chemotherapy (Rao et al., 2017). In addition, knowledge of the fundamental principles for formation of natural extracellular glycoprotein networks will guide the design of artificial scaffolds for tissue engineering and other biotechnological applications.

Studying large and flexible glycoproteins is challenging. Various models for the mucin and VWF amino-terminal regions have been put forth based on low-resolution structural data (Huang et al., 2008; Ambort et al., 2012; Nilsson et al., 2014; Ridley et al., 2014; Trillo-Muyo et al., 2018), but a consistent picture for their arrangement and assembly mechanism had yet to emerge. An X-ray crystal structure of a monomeric version of the ∼400 amino-acid VWF D3 segment provided important information on the protein folds that make up the mucin and VWF multimerization regions (Dong et al., 2019). We recently reported the high-resolution structure of a dimer of the D3 segment of the intestinal gel-forming mucin MUC2 showing the intermolecular disulfide bonding pattern (Javitt et al., 2019). However, describing how this domain fits into the temporal and physical scheme of hydrogel construction required a broader context, which we provide by the study of larger functional fragments of mucins.

## Results

### MUC2 forms a beaded polymer at acidic pH

To determine the mechanism of mucin multimerization, we produced and analyzed the entire ∼1400 amino-acid amino-terminal region of MUC2. This region, designated D1D2D3CysD1, comprises three “D” segments, a proline-, threonine-, and serine-rich stretch (PTS) of 40 amino acids, and a CysD domain (Figure 1C). A homologous region is found in the respiratory mucins MUC5B and MUC5AC, as well as in VWF (Figure S1), except that the latter lacks a PTS segment and CysD domain.

We observed that MUC2 D1D2D3CysD1 undergoes robust but reversible polymerization at acidic pH, forming long filaments composed of distinctive “beads” as visualized by electron microscopy (EM) (Figure 2A). A MUC2 fragment lacking CysD1 did not polymerize under similar conditions (not shown), whereas the D1D2D3CysD1 portion of murine respiratory Muc5b formed longer filaments than those previously observed for a human D1D2D3 MUC5B fragment (*i.e*., lacking CysD1) (Figure 2A) (Trillo-Muyo et al., 2018). High-throughput dynamic light scattering showed that MUC2 D1D2D3CysD1 polymerization proceeded in a reproducible and controlled manner between pH 5.4 and pH 6.2 at 37 °C (Figure 2B). Though closed loops were occasionally seen for our His_6_-tagged protein (Figure 2A), we did not observe a preference for the hexagonal structures reported previously for a Myc- and EGFP-tagged version of a similar MUC2 fragment (Ambort et al., 2012: Nilsson et al., 2014). Consistent with amino-terminal dimerization rather than the proposed trimerization of native MUC2, cleavage of O-glycan-rich regions with the protease StcE (Malaker et al., 2019) to liberate mucin globular domains yielded a product of similar size for mouse colon Muc2 as for recombinant MUC2 D1D2D3CysD1 (Figure 2C), which was shown to be a disulfide-bonded dimer (Javitt et al., 2019). StcE-treated murine (Figure 2C) and human (Figure 2D) fecal mucins were detected as migrating similarly to disulfide-bonded recombinant D3 dimers. In sum, these results demonstrate that endogenous and recombinant gut mucins have similar multimerization properties, consistent with amino-terminal disulfide bonded dimerization as the shared mode of assembly for both gut and lung gel-forming mucins.

**Figure 2.**
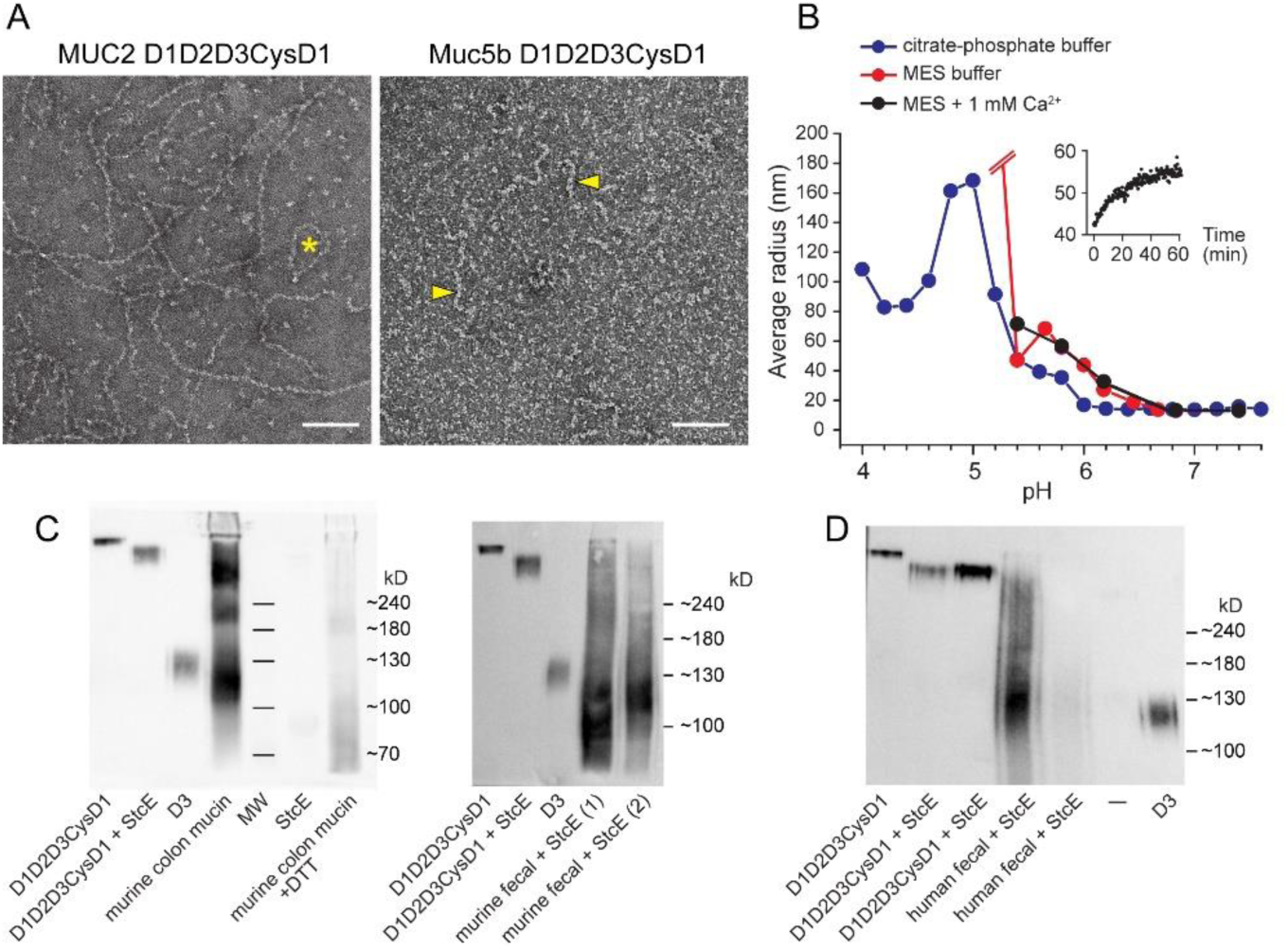
Polymerization of the mucin amino-terminal region. (A) Polymerization of the D1D2D3CysD1 segments of MUC2 and Muc5b upon incubation with 10 mM CaCl_2_ at pH 6.2 and pH 5.4, respectively, visualized by negative stain EM. Scale bars are 100 nm. The asterisk indicates a looped MUC2 polymer, and the arrowheads point out Muc5b polymers. (B) Dynamic light scattering analysis of MUC2 D1D2D3CysD1 as a function of pH. Average radius (averaged after the signal plateaued) corresponds to the size of a sphere with similar diffusion. The sample in MES buffer at pH 5.1 gave a calculated radius of about 300 nm. The inset shows a sample trace (MES pH 5.8) of polymerization as a function of time. (C) Western blot of murine colon and fecal mucus digested with StcE protease compared to recombinant MUC2 fragments. The Muc2 fragments corresponding to D1D2D3 and D3 migrate slightly faster than their human recombinant counterparts. Detection was done using an antibody raised against the murine Muc2 D3 region. The disappearance of bands upon reduction with dithiothreitol (DTT) is due both to disassembly of disulfide-bonded dimers and to loss of conformational epitopes. The two fecal samples show the variability of Muc2 degradation products. (D) Western blot of human fecal mucus digested with StcE protease compared to recombinant MUC2 fragments. Detection was done using an antibody raised against the human Muc2 D3 region. Two concentrations of StcE-digested recombinant D1D2D3CysD1 were applied to the gel.

### Structure and domain organization of the MUC2 multimerization region

Single particle cryo-EM analysis and 3D reconstruction were performed on polymerized MUC2 D1D2D3CysD1 vitrified after exposure to pH 5.7. Polymers were segmented using an automated particle picking algorithm, and boxed segments were treated as single particles. After image classification and refinement, a density map at an overall resolution of 2.95 Å was produced (Figure S2, Table S1). The map corresponded to an individual bead in the D1D2D3CysD1 polymer and contained the contacts to the adjacent beads. The model built into this map showed that successive beads were related by a rotation of 98.5° around the polymer axis and displaced by 13.7 nm (Figure 3A, Supplementary movie 1). It is important to emphasize that the polymers described herein are not the expanded head-to-head and tail-to-tail strands of full-length mucins but rather represent the compact assembly state during formation and storage of polymers at low pH in the Golgi apparatus and secretory granules. Analysis of the compact polymers as performed in this work illuminates the mechanism for biosynthesis of full-length mucin polymers.

**Figure 3.**
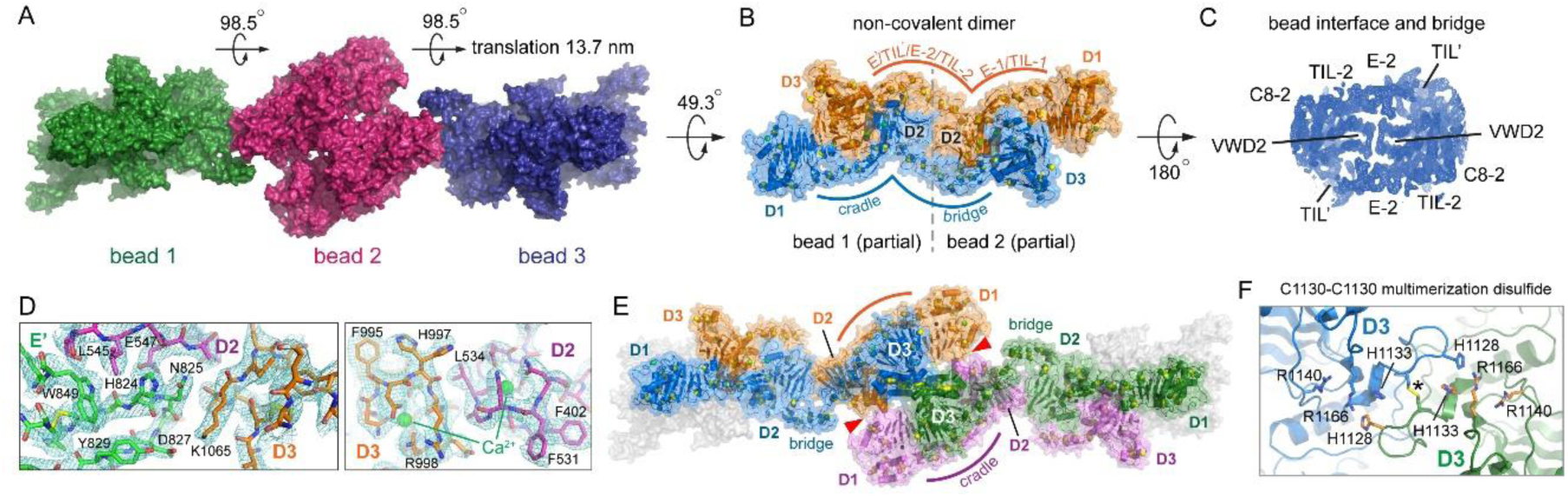
Structure of the MUC2 amino-terminal polymer. (A) Three MUC2 beads are shown with rotations and translations indicated. (B) Two D1D2D3CysD1 molecules, colored orange and blue, show how MUC2 polypeptides span multiple beads. Disulfide bonds are indicated as spheres with yellow sulfurs. The “cradle” is the cysteine-rich spacer between D1 and D2 that accommodates the D3 domain from the second subunit of the non-covalent dimer. The “bridge” is the cysteine-rich spacer between D2 and D3 that enables D3 to extend into the next bead. The domain composition of the cradle and bridge are indicated above the orange molecule. (C) Cryo-EM map of the inter-bead junction demonstrating that the cysteine-rich bridge TIL-2/E-2/TIL’/E’ crosses from one bead to the next. The E’ domain is below density corresponding to other portions of the molecule and is not labeled. (D) Cryo-EM density map and structure of selected interface regions between D3 and the second subunit in the non-covalent dimer. (E) The bead structure is formed by two noncovalent dimers (dimer 1: blue and orange; dimer 2: magenta and green) linked by disulfide bonds through a D3 domain contributed by each dimer (green and blue). Red arrowheads indicate the D1-D2 contacts between the two dimers. (F) Clusters of histidine and arginine side chains straddle the intersubunit disulfide (indicated by *) between Cys1130 and its symmetry mate.

From the cryo-EM density map, most of the D1D2D3CysD1 amino acid sequence could be readily traced and the abundant disulfides assigned (Table S2). The MUC2 beads make contacts along the polymer by edgewise interaction of apposed D2 β-sandwiches (Figure 3B). Additionally, contiguous density was found circumscribing the β-sandwiches and reaching from one bead to the next, as confirmed by a reconstruction performed using a larger box size to improve the map at the inter-bead interface (Figure 3C). This density demonstrates that a single MUC2 molecule exceeds the boundaries of an individual bead and projects its D3 domain into the adjacent bead (Figure 3B). The adjacent bead in exchange donates a different D3 domain to the original bead. This reciprocal arrangement, which was not anticipated by previous low-resolution models for mucins or VWF (Huang et al., 2008; Trillo-Muyo et al., 2018), is enabled by the TIL and E modules (Figure 1C). These disulfide-rich modules serve as spacers between the VWD domains: the TIL/E region between D1 and D2 forms a “cradle” for D3 of another protein molecule, while the duplicated TIL/E region between D2 and D3 provides the necessary length to bridge between beads (Figure 3B). Contacts between D3 and the cradle include conserved histidines and other polar and charged amino acids (Figure 3D), and the D2/D3 interaction juxtaposes two calcium binding sites (Figure 3D), consistent with pH- and calcium-dependent interactions. A major conclusion from the MUC2 structure is that the disulfide-rich linker TIL/E modules hold apart the D3 domains of the two MUC2 molecules within an extended dimer, providing a mechanism that forbids disulfide bonding between D3 domains at the first step of assembly and preserves them available for subsequent events.

### Disulfide-mediated MUC2 polymerization

In addition to showing how MUC2 and related molecules evolved to block disulfide bonding between the two D3 domains within the same intertwined dimer, the cryo-EM structure also revealed how MUC2 self-assembly facilitates disulfide bonding between D3 domains from separate dimers. Contacts are formed in the beaded polymer between D1 and D2 of two non-covalent dimers in a manner compatible with the apposition of the D3 domains (Figure 3E). Within this arrangement, D1/D2 cradles D3 with cysteines Cys1088 and Cys1130 accessible (Figure S3) for their participation in the intermolecular disulfide bonds first seen in the crystal structure of the isolated D3 domain (Javitt et al., 2019). In the crystals, which were grown at pH 4.2, histidine side chains near the Cys1130-Cys1130 disulfide were disordered. In the cryo-EM structure, of a sample prepared at pH 5.7, these histidines participate in symmetry-related histidine/arginine clusters formed across the D3 dimer interface (Figure 3F). These clusters may determine the pH range at which the D3-D3 interface forms and control the disulfide-mediated polymerization step. The process of bringing non-covalent dimers together to form the bead, which is expected to occur in the second stage of mucin and VWF assembly and to be tunable by pH, juxtaposes the appropriate D3 domains for generation of disulfide-linked polymers.

### Model of VWF tubule assembly

Though many aspects of domain structure and assembly mechanism are shared between mucins and VWF, VWF uniquely forms tubules prior to secretion. Low-resolution reconstructions of these tubules (Huang et al., 2008; Berriman et al., 2009) have provided an initial glimpse of their organization, but, without assignment of domain identity and connectivity, it was unclear how tubules relate to formation of VWF covalent polymers. The homologous MUC2 bead structure appeared compatible in size and organization with the repeat unit in the VWF tubule, allowing MUC2 to illuminate VWF assembly. By comparison with the MUC2 bead reconstruction, the ambiguity in domain positions in VWF (Huang et al., 2008) was resolved, and the connectivity between domains was corrected (Figure 4A).

**Fig. 4.**
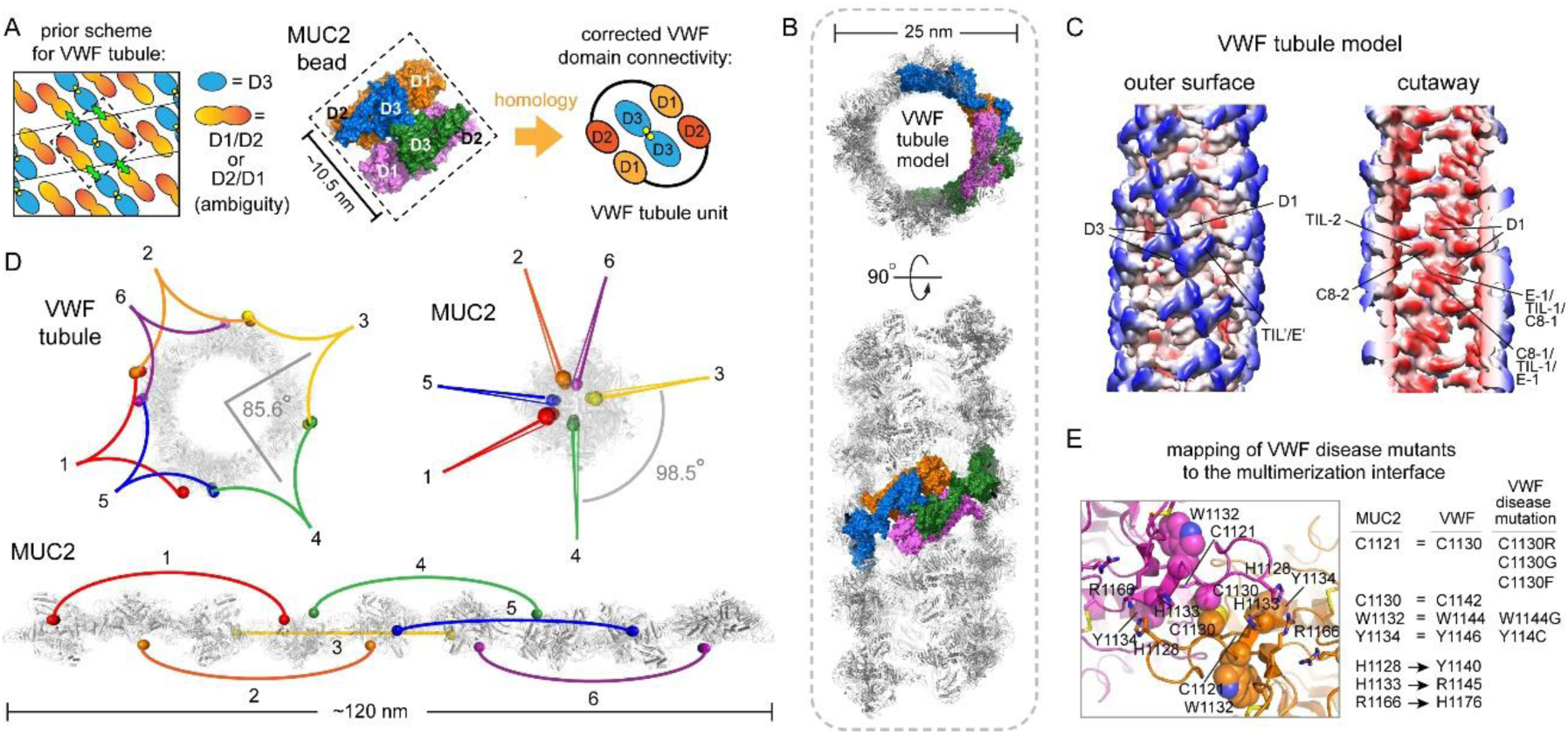
Comparison of MUC2 with the VWF tubule. (A) The MUC2 bead structure resolves ambiguities in a VWF tubule model based on a low-resolution helical reconstruction (Huang et al., 2008). (B) Homology model of the VWF tubule. Four polypeptide chains are colored to correspond to Figure 3E. (C) The VWF tubule homology model was downsampled to 6 Å resolution for comparison with previous low-resolution studies (Huang et al., 2008). Coloring is red to blue according to distance from the tubule axis. Features of the tubule that can now be unambiguously assigned are labeled. (D) Views down the tubule and polymer axes show how carboxy-terminal regions of VWF and MUC2 would spiral around the assemblies. The carboxy termini of the VWF D3 segments (residue 1207) are labeled with spheres colored to indicate the subunits expected to be paired *via* the carboxy-terminal regions (not present in this study). Numbering indicates successive carboxy-terminal disulfide-linked partners. For MUC2, the last amino acid from the cryo-EM structure (residue 1391) is similarly indicated by colored spheres. The carboxy-terminal regions of MUC2 are expected to spiral around the amino-terminal polymer in a manner analogous to the VWF tubule. Below is a side view of the MUC2 polymer. (E) Conserved amino acids near the MUC2 dimer interface are mutated in VWF disease. Side chain atoms of disease mutation positions (de Jong and Eikenboom, 2017) and the C1130 disulfide are shown as spheres. Amino acid numbering of corresponding residues in MUC2 and VWF are indicated, and VWF disease mutations in this region are listed.

The homology with MUC2 was further exploited to generate a detailed model for the VWF tubule using the known rotation and translation parameters (Huang et al., 2008) (Figure 4B). Importantly, an insertion of about 20 residues in the linker between D2 and D3 of VWF (Figure S1), which contains a furin protease cleavage site (Springer, 2014), also accommodates the different orientation of beads in the VWF tubule compared to the MUC2 polymer. The VWF tubule model based on the MUC2 bead recapitulates the features of the low-resolution VWF reconstruction and additionally reveals the positions of the domains and cysteine-rich linkers (Figure 4C). Lateral contacts contributing to tubule formation are made by reciprocal interactions between the disulfide-rich segments corresponding to C8-1, TIL-1, and E-1 of MUC2 (VWF residues 215-374), which constitute the outer edge of the cradle. The D3 domains are on the outer face of the tubule. Consistent with previous suggestions (Huang et al., 2008), successive disulfide-linked pairs of VWF carboxy-terminal regions would spiral around the outside of the tubule at ∼85° intervals due to the rotation angle between repeat units (Figure 4D). Based on the orientation of beads in the MUC2 polymer, successive disulfide-linked pairs of MUC2 PTS and carboxy-terminal regions are predicted to spiral around the MUC2 amino-terminal region polymer in ∼98° steps (Figure 4D). Though MUC2 does not form tubules like VWF, the MUC2 beaded polymer can be considered a degenerate helix with an approximate (slightly over-wound) four-fold screw, whereas the VWF tubule has 4.2 subunits per helical turn. In both cases, this angular separation of successive units allows the remaining portions of VWF and mucins, which for MUC2 equates to multiple megadaltons of glycoprotein per subunit, to project in all directions from the central spine of the amino-terminal assembly.

### CysD domains and O-glycosylated PTS region

VWF assembly is stabilized by interactions both along the helical polymer and between successive turns of the helix. MUC2 appears to lack the latter contacts and thus depends on other features to stabilize the beaded polymer. As noted above, polymerization of the MUC2 amino-terminal region (Figure 2A) is not detected in the absence of the CysD1 domain, so the important role of this domain remains to be described. CysD1 spans the last ∼100 amino acids of the MUC2 fragment (Figure 1C). The CysD domain fold was not known prior to this study, so the CysD1 structure was determined at a higher resolution (1.63 Å) by X-ray crystallography (Table S3), in parallel with its visualization by cryo-EM within the MUC2 amino-terminal bead. The connectivity of cysteines in CysD1 disulfides (Figure 5A) was found to contradict the arrangement proposed previously for the homologous MUC2 CysD2 domain (Ambort et al., 2011). The CysD1 domain consists of a small β-sandwich with three β-strands per β-sheet and two bound calcium ions (Figure 5B, C). A characteristic feature of mucin CysD domains is the presence of a Trp-X-X-Trp (WXXW) motif similar to the C-mannosylation motif identified in proteins with thrombospondin type I repeats (Niwa and Simizu, 2018). Consistent with observations for CysD2 (Ambort et al., 2011), the tryptophans in the recombinant proteins used for cryo-EM and crystallographic structure determination were not mannosylated, but the C2 carbons of the tryptophan indole rings are exposed in the structures and thus are sterically accessible for modification. In fact, the geometry of the WXXW motif and its context within a Trp-Arg/Lys ladder are very similar to the analogous motif in thrombospondin, despite the substantial difference between the folds (Figure 5D).

**Figure 5.**
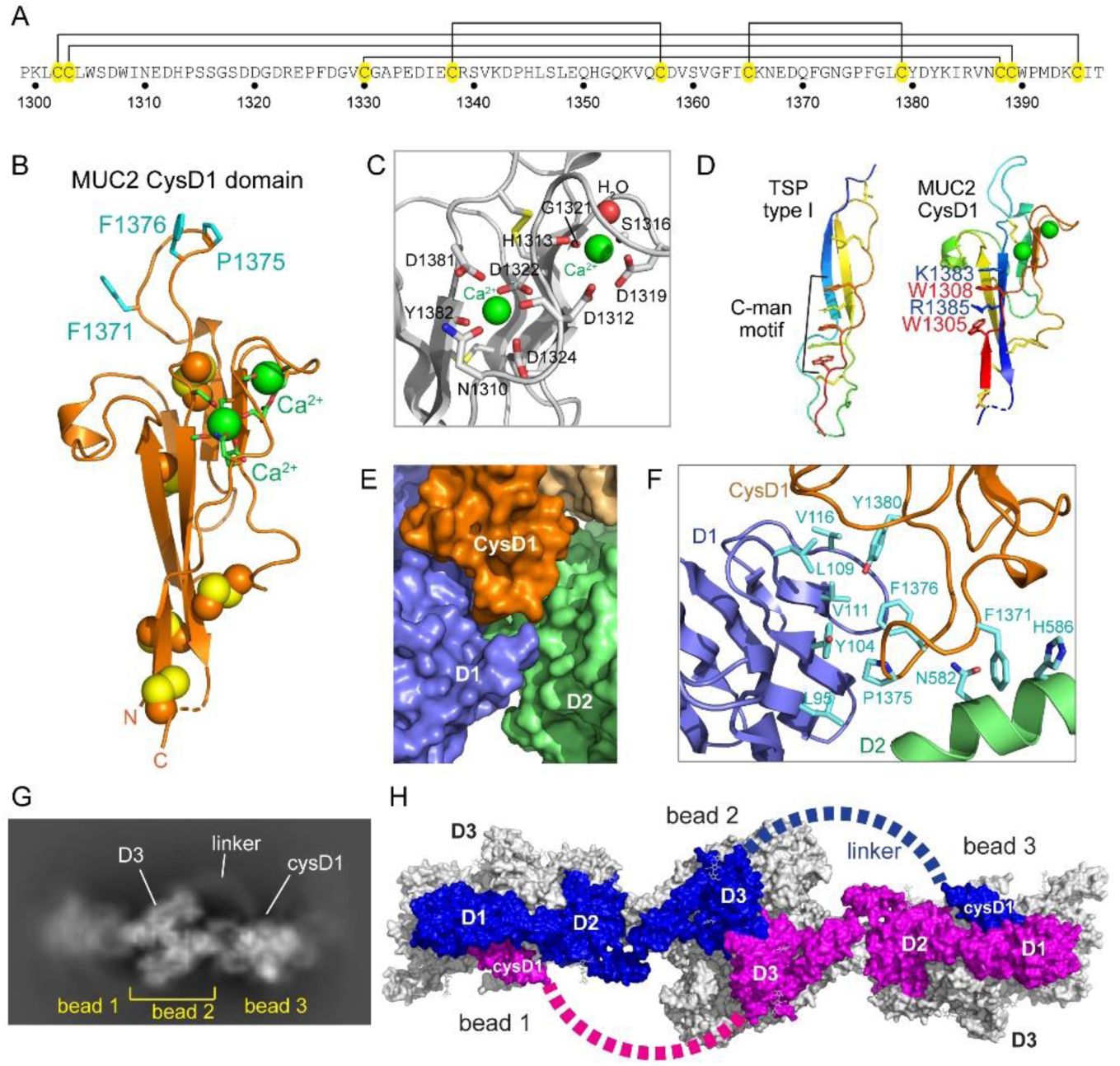
CysD1 domain structure and O-glycosylated PTS linker. (A) Disulfide map of MUC2 CysD1. Amino acid numbering corresponds to the full-length MUC2 sequence. (B) Ribbon diagram of the MUC2 CysD1 domain. Green spheres are bound calcium ions; other spheres are disulfide bonded cysteines. Exposed hydrophobic side chains involved in intermolecular contacts are shown in cyan. (C) Calcium coordination in the CysD1 domain. (D) Comparison of the tryptophan-arginine/lysine ladder in thrombospondin (TSP) type I repeats (PDB ID 1LSL) and MUC2 CysD1. (E) Binding by CysD1 reinforces the intermolecular interactions between D1 and D2 of the non-covalent dimers making up the MUC2 bead. (F) Contacts at the D1/D2/CysD1 interface. (G) An arc of density in a representative 2D class of a two-bead reconstruction links a D3 domain with a CysD1 domain in the adjacent bead. (H) With the contribution of the CysD1 domain, each MUC2 polypeptide chain spans three beads.

Within the context of the MUC2 polymer, CysD1 was seen to reinforce the D1-D2 dimer-dimer contacts and thereby support the bead structure (Figure 5E). It does so by clasping the edge of one sheet of the VWD1 β-sandwich, making a number of hydrophobic contacts while also binding the D2 domain of the second dimer in the bead through a potentially pH-sensitive cation-π interaction between Phe1371 and His586 (Figure 5F). These contacts explain the requirement for the CysD1 domain in MUC2 polymerization.

Importantly, the D3 and CysD1 domains are on opposite sides of an individual MUC2 D1D2D3CysD1 bead. The intervening primary structure includes a 40-amino acid PTS segment (Figure 1C), which could potentially circle the bead to link D3 with CysD1. However, the data suggest a different solution: an arc of density connecting D3 with the CysD1 domain of the adjacent bead was seen in the cryo-EM 2D classes (Figure 5G) and in the unfiltered 3D reconstruction (Supplementary movie 2). When CysD1 is considered, each MUC2 polypeptide thus extends over three beads in the polymer (Figure 5H), and each dimer linked by carboxy-terminal CTCK disulfides (Zhou et al., 2014) would extend over four beads (Figure 4D). The visibility of the inter-bead arc of density also suggests that the PTS region is not completely disordered but rather displays some regularity, in line with a recent study of longer O-glycosylated mucin regions (Hughes et al., 2019). Notably, the PTS linker and the observed CysD1 interactions would preclude a tubular arrangement for mucins as later evolved for VWF, since they would position the rest of the mucin molecule within the central cavity of the tubule. Without the tight helical arrangement of VWF tubules, mucins avoid the topological problem of where to pack the mass of glycoprotein that exists carboxy terminally to CysD1.

## DISCUSSION

The cryo-EM structure of MUC2 D1D2D3CysD1 and the analysis presented above lead to a unified assembly model explaining the origin of head-to-head and tail-to-tail polymers (Figure 6). Upon biosynthesis (Figure 6A), two protein molecules dimerize non-covalently by a reciprocal exchange of D3 domains (Figure 6B), which dock into cradles formed by the D1/D2 domains and intervening cysteine-rich spacers (Figure 3B). It is presumed that these same two molecules also become disulfide linked through the CTCK domains at their carboxy termini (Zhou et al., 2014) (Figure 6B). D3 domains are thereby isolated from one another in the CTCK-linked dimer, such that formation of a closed dimer (disulfide bonded at both ends) is prevented. Later in the secretory pathway, these dimers become disulfide bonded to other dimers *via* their D3 domains (Figure 6C), promoted by the pH gradient along the secretory pathway. Overall, this multi-step process forms the linear mucin beaded polymer described in this work and produces the helical tubules of VWF. For both assemblies, the remainder of the glycoprotein would project outwards (Figure 4D). Despite their apparent differences, fundamental topological principles are shared by mucins and VWF: the bead rotation along the polymer axis in mucins is replaced by the bead rotation around the tubule axis in VWF. Release of non-covalent interactions or proteolytic cleavage of D1/D2 produces the final stretched polymers that function extracellularly *in vivo* (Figure 6D).

**Figure 6.**
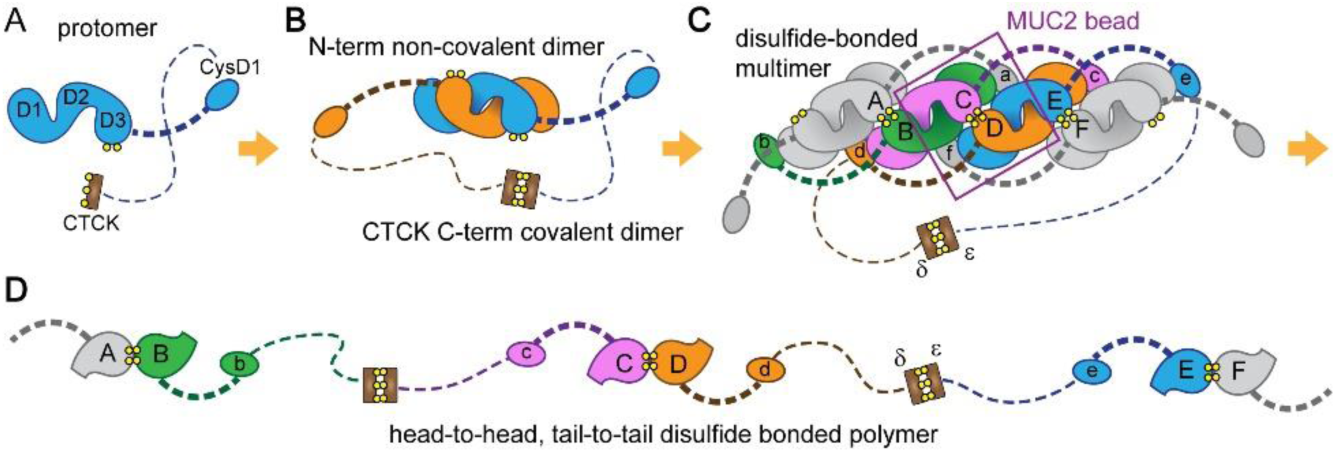
Stepwise assembly mechanism for mucins. (A) The TIL/E spacers ensure the physical separation of the D1, D2, and D3 domains upon folding of the MUC2 protomer. Dashed lines correspond to PTS regions (not drawn to scale); thicker dashed lines represent the 40 amino-acid segment between D3 and CysD1, while thinner dashed lines represent the thousands of amino acids between CysD1 and the CTCK domain (other carboxy-terminal domains, recently studied structurally [Ridley et al., 2019] are not represented). (B) Two protomers form a non-covalent dimer *via* interactions between D3 domains and the cradle formed by D1/D2. The CTCK region forms a disulfide-bonded dimer. (C) Pairs of dimers further assemble into a compact polymer, promoting disulfide bond formation between D3 domains. D3, CysD1, and CTCK domains from the same molecule are annotated by corresponding letters (*e.g*., D, d, δ). One bead within the compact polymer (boxed) contains domains from six distinct molecules: A in gray (CysD1 domain), B in green (D1 and D2 regions), C in magenta (D3 region), D in orange (D3 region), E in blue (D1 and D2 regions), and F in gray (CysD1 domain) (Supplementary movie 2). (D) Upon secretion, the polymer may be extended by release of the D3 domains from the D1/D2 cradle after breaking non-covalent interactions or by proteolytic removal of D1/D2. The end result is a head-to-head, tail-to-tail disulfide bonded polymer. The essence of the assembly model also applies to VWF.

Particularly intriguing elements specific to mucin assembly are the CysD modules. MUC2 has only two CysD domains, whereas lung mucins have many more, interspersed with PTS regions (Figure 1C). The interaction mode observed for MUC2 CysD1 may be generally informative regarding CysD contacts in mucins. It has been reported that an isopeptide bond forms between the MUC2 D2 domain and CysD2, which is separated from CysD1 by about 400 amino acids of PTS region (Figure 1C) (Recktenwald and Hansson, 2016). Isopeptide bonds may be an extracellular post-translational mucin modification that helps form an impenetrable cross-linked network once linear polymers are secreted. Supporting a general mode for CysD function, an interaction between CysD2 and D2 with similar geometry as that reported here between CysD1 and D1 would juxtapose the isopeptide-linked amino acids, Gln470 from D2 and Lys1796 from CysD2. Though Gln470 is buried in the low-pH MUC2 polymer by the inter-bead bridge, it may be exposed after polymer expansion as the pH increases during secretion into the gut lumen. Another intriguing observation is that transgenic mice expressing artificial stretches of tandem CysD domains showed increased robustness of mucin gels and improved intestinal barrier function (Gouyer et al., 2015). Knowledge of the positions and roles of CysD domains in mucin multimerization will now guide the design of CysD constructs with improved composition and spacing for desired physiological effects.

Due to the recalcitrance of mucin PTS regions to recombinant production and high-resolution structural study, the mechanism described herein focuses on the disulfide-rich and globular amino-terminal portion of gel-forming mucin molecules. However, supramolecular assembly *in vivo* accommodates the massive full-length glycoproteins. The supramolecular architecture of intact VWF is reflected in the morphology of the intracellular compartment in which VWF is stored prior to secretion, and a similar comparison between mucin molecular structure and its storage compartments may shed light on mucin assembly in cells. The VWF-packed Weibel-Palade bodies in endothelial cells are elongated vesicles with a diameter of about 0.15 µm and lengths of a few micrometers, containing bundles of about nine tubules (Huang et al., 2008; Berriman et al., 2009). In contrast, MUC2 granules are much larger, roughly spherical structures about 10 µm in diameter. Each of these large mucin granules comprises numerous sub-compartments, which originate from distinct membrane-delimited mucin-containing vesicles. The apparent isotropy of mucin vesicles can be reconciled with the linear MUC2 amino-terminal polymers observed and studied in this work, as the polymers are likely to have only short-range order and not to be aligned with neighboring polymers within the granule. The cryo-EM micrographs used for the MUC2 bead reconstruction show kinks and curvature in the beaded chains, with regular polymers rarely exceeding ten beads, corresponding to about 0.14 µm (Figure S2D). To enable comparison of MUC2 condensed-polymer persistence lengths with granule dimensions, we quantified the sub-compartment areas from a large number of micrographs of murine colon sections. From these data, it was determined that the sub-compartments would have an average diameter and volume of 1.3 µm and 1.2 µm^3^, respectively. These measurements indicate that the mucin vesicle dimensions are nearly an order of magnitude larger than the scale on which regularity was observed for the recombinant MUC2 polymers. Therefore, our observations are consistent with previous suggestions that intracellular condensed mucins have locally ordered domains but lack an overall directionality on the micron-length scale and undergo isotropic swelling upon exocytosis (Viney et al., 1993). It remains to be determined whether the long PTS segments and carboxy-terminal mucin regions affect the properties and persistence length of condensed polymers formed by intact glycoproteins packaged *in vivo*.

An important discovery made during structural analysis of MUC2 is that the lengths of PTS segments in mucins can have particular significance in determining the range of potential interactions made by CysD domains. We observe that the ∼40-amino acid PTS segment downstream of the D3 domain allows the tethered CysD1 domain to bind the neighboring bead in the polymer. Presumably, a shorter PTS region would not reach, whereas a longer PTS region would overshoot if relatively rigid or, if flexible, would introduce a higher entropic barrier to fixing the position of the CysD1 domain. The lengths of the PTS regions vary between different pairs of CysD domains, between mucin paralogs, and between species. Mucin PTS length polymorphisms also exist within the human population and have been identified as modifiers of disease severity in cystic fibrosis (Guo et al., 2011). These differences suggest that mucins have evolved their great diversity of viscoelastic properties through tuning the number and geometry of interactions involving CysD domains attached to PTS tethers. The evolvability of CysD/PTS interactions thus complements the conservation of the core mucin/VWF assembly pathway revealed in this work and expands mucin functionality. The fundamental principles of mucin architecture described here will provide focus for the development of drugs and methods to enhance and restore the protective coatings of epithelial and endothelial tissues.

## Data availability

Atomic models were deposited to the protein data bank under accession codes 6TM6 (CysD1) and 6TM2 (D1D2D3CysD1). The EM density map was deposited to the EMDB under accession code 10517.

## Acknowledgements

The authors acknowledge Karin Strijbis and Emmanuel Levy for helpful suggestions. This work was supported by the European Research Council (grant number 310649), by the I-CORE Program of the Planning and Budgeting Committee and the Israel Science Foundation (grant No. 1775/12), and by a research grant from the Center for Scientific Excellence at the Weizmann Institute of Science.

## Author contributions

G. J. and D. F. conceptualized the research. G. J. and N. E. collected cryo-EM data. G. J. performed single-molecule reconstruction with the help of N. E. and R. D. Structure modeling into the cryo-EM map was performed by D. F. together with G. J. Crystallization, data collection, and structure solution were performed by L. K., with help from D. F. Together with G. J., L. A. contributed to protein purification and dynamic light scattering.

D. F. and G. J. wrote the manuscript with contributions from all other authors.

## Supplemental Information

### Materials and Methods

#### Protein production and purification

D1D2D3CysD1 and D1D2D3 were produced by transient transfection of HEK293F cells as described (Javitt et al., 2019). To produce the CysD1 domain in isolation, residues 1299 to 1397 of human MUC2 were inserted into the pcDNA3.1 plasmid downstream of a segment encoding the sequence MRRCNSGSGPPPSLLLLLLWLLAVPGANAAPQGHHHHHHENLYFQGG, which includes the signal sequence from the protein QSOX1, a His_6_ tag for purification, and a TEV cleavage site for removing the tag. TEV cleavage leaves two non-native glycines fused to the amino terminus of the CysD1 domain. Transfection and nickel-nitrilotriacetic acid (Ni-NTA) chromatography were performed as described (Javitt et al., 2019). The eluted protein was dialyzed against 0.5X phosphate buffered saline at room temperature overnight at a 25:1 molar ratio with TEV protease. After dialysis, Ni-NTA beads were added to the solution, and the suspension was mixed end-over-end for 20 min. The beads were then allowed to settle to remove TEV protease, the cleaved His_6_ tag, and any remaining uncleaved material. The supernatant was collected, the cleaved protein was concentrated to 7 mg/ml, and the buffer was exchanged to 10 mM Tris, pH 7.5, 20 mM NaCl for crystallization.

#### Dynamic light scattering

Samples of purified recombinant D1D2D3CysD1 were placed in a clear bottom black 96 well plate and diluted to 0.3 mg/ml in 37 °C pre-warmed 180 µM citrate-phosphate or MES buffer with 150 mM NaCl, with or without 1 mM CaCl_2_. Dynamic light scattering was recorded using a DynaPro Plate reader (WYATT Technology) pre-warmed to 37 °C. The dead time at the beginning of each experiment due to plate insertion and temperature re-equilibration was about 6 min. Average radius was calculated by averaging the signal between 40 and 120 min after the start of the experiment. Data were processed with the supplied Dynamics software (Wyatt Technology).

#### Colon mucus digestion and western blotting

StcE enzyme was produced as described (Yu et al., 2012) and exchanged into phosphate buffered saline (PBS). A 2 cm-long section of murine distal colon (strain C57bl, age 6 months) was cut longitudinally to expose the luminal surface, onto which 100 µl of 10 µM StcE, diluted 10X into water from a 100 µM stock in PBS, was pipetted. The colon section was placed into a microcentrifuge tube containing an additional 200 µl of diluted StcE and incubated for 3 hours at 37 °C. Toward the end of this incubation, StcE was added at a concentration of 1 µM to 1.5 µM D1D2D3CysD1 in a microcentrifuge tube and incubated for 0.5 hour at 37 °C. The StcE solution was then pipetted gently up and down on the colon section to suspend digested mucus material, which was transferred to a microcentrifuge tube. Seventy-five µl fresh PBS was pipetted onto the colon section, mixed to release any remaining mucus, and then added to the material in the tube. The extracted digested mucus was spun at high speed in a microfuge for 1 min to pellet insoluble material. Ten microliters of the soluble fraction was applied without reduction to a 7.5% polyacrylamide gel alongside 10 µl each of 1.5 nM D3, undigested D1D2D3CysD1, and StcE-digested D1D2D3CysD1, after mixing 1:1 with Laemmli buffer. Proteins were transferred to nitrocellulose for western blotting, and detection was done using an antibody raised against the murine Muc2 D3 region. For fecal mucin samples, 20 mg feces was mixed into 150 µl of 10 µM StcE and incubated for 3 hours at 37 °C, with periodic vortexing. At the end of the incubation the samples were spun at high speed in a microfuge for 1 min to remove insoluble material. Recombinant samples were prepared as above. The western blot of human fecal mucus was detected using an antibody raised against the human MUC2 D3 region.

#### Sample preparation for EM

For negative staining, purified D1D2D3CysD1 protein was incubated at a concentration of 1 mg/ml in 50 mM MES, pH 5.4 to 6.2, 225 mM NaCl, with or without CaCl_2_ at 37 °C for 24 hours. The protein was then diluted to a concentration of 0.03 mg/ml, and 3 µl solution was applied to a glow discharged carbon-coated 300 mesh copper grid (Electron Microscopy Sciences) for 30 seconds, followed by staining with 2% uranyl acetate solution. Samples were visualized using a Tecnai T12 electron microscope (Thermo Fisher Scientific) equipped with a OneView camera (Gatan). For CryoEM, purified D1D2D3CysD1 protein was incubated at a concentration of 0.3 mg/ml in 50 mM MES, pH 5.7, 225 mM NaCl at 37 °C for 24 hours. The incubated protein solution (3 µl) was pipetted onto glow discharged Quantifoil R1/2, 300 mesh copper grids. Grids were plunge frozen into liquid ethane cooled by liquid nitrogen using a Vitrobot plunger (Thermo Fisher Scientific) at 100% humidity.

#### Cryo-EM image acquisition

Cryo-EM data were collected on a Titan Krios G3i transmission electron microscope (Thermo Fisher Scientific) operated at 300 kV. Movies were recorded on a K3 direct detector (Gatan) installed behind a BioQuantum energy filter (Gatan) using a slit of 20 eV. Movies were recorded in counting mode at a nominal magnification of 105,000x, corresponding to a physical pixel size of 0.85 Å. The dose rate was set to 23 e^-^/pixel/s, and the total exposure time was 1.5 s, resulting in an accumulated dose of ∼48 e^-^/Å^2^. Each movie was split into 45 frames of 0.033 s. Nominal defocus range was -1 to -2 μm. SerialEM was used for automated data collection (Mastronarde, 2005), in which a single image was collected from the center of each hole. Image shift was used to navigate within 3×3 hole arrays, and stage shift to move between arrays. Beam tilt was adjusted to achieve coma-free alignment when applying image shift.

#### Cryo-EM image processing and atomic coordinate fitting

Image processing was performed using CryoSPARC software v2.9 (Punjani et al., 2017). A total of 3242 acquired movies were subjected to patch motion correction, followed by patch CTF estimation. Of these, 2745 images showing CTF fit resolution better than 3.5 Å were selected for further processing. Initial particle picking was done using the Blob Picker function on a subset of 200 micrographs. About 32,000 particles were extracted, 2D classified into 50 classes, and 15 class averages with high resolution and visible secondary structure were used as templates for automated particle picking from the entire data set. In total 987,943 particles were extracted and subjected to multiple rounds of 2D classification, in which 2D classes were selected for further processing based on the appearance of secondary structure elements, resulting in a dataset of 84,970 particles. These particles were used for *ab initio* 3D reconstruction, followed by homogeneous 3D refinement, which resulted in a refined map at 3.2 Å resolution. To extract more and better particles from the micrographs, 50 equally spaced re-projections of the new reconstruction were used as templates for another round of automated particle picking, resulting in 1,430,962 picked particles. The new data set was, similarly, subjected to multiple rounds of 2D classifications, retaining classes with well-resolved features, and 293,080 particles were retained and used for 3D refinement. Subsequently, 3D heterogeneous refinement into 3 classes was performed, and one class, consisting of 178,136 particles, which showed the highest resolution, was homogeneously refined with C2 symmetry to 2.95 Å resolution. The final map was subjected to local resolution estimate and filtered accordingly. To generate the two-bead reconstruction, these particles were re-extracted with a larger box size and subjected to homogeneous refinement to 3.95Å resolution with C1 symmetry.

#### Model building, refinement, and analysis

The previously reported MUC2 D3 dimer (PDB ID 6RBF) was fitted manually into the single-bead cryo-EM map, and the rest of the model was manually built in Coot (Elmsley et al., 2010). Each β-sandwich VWD and helical-bundle C8 subdomain of regions D1 (residues 35-291; numbering according to Uniprot Q02817, domain boundaries are according to ref. 5), D2 (residues 388-655), and D3 (residues 858-1117) (Fig. 1C) were readily built into the structure model. In addition, density was evident for the TIL-1 and E-1 subdomains (residues 292-387) linking D1 to D2, as well as for most of TIL-2 (residues 656-719) and E’ (residues 822-857), which contribute to bridging D2 and D3. Partial density was also observed for the remainder of the D2-D3 bridge formed by E-2 and TIL’. The position of the CysD1 domain was clear in the map. The trajectory of the PTS region linking D3 with CysD1 could be seen, but this region could not be reliably modeled. The final model refined against the 2.95 Å map included MUC2 amino acids 35 to 721, 724 to 749, 780 to 793, 801 to 1197, and 1302 to 1391. The model was refined by cycles of real-space refinement using Phenix (Adams et al., 2010) and manual rebuilding in Coot. The structure was analyzed and structure figures were generated using pymol (DeLano, 2002). Model geometry was assessed using Molprobity (Chen et al., 2010). The entire spacer between D2 and D3 was traced and the disulfide connectivity determined in the context of the two-bead map.

#### CysD1 crystallization

CysD1 plate-like crystals appeared by the hanging drop vapor diffusion method over a well solution containing 100 mM citrate/phosphate buffer, pH 4.4, with 15% ethanol and 1% PEG 1K within 12 hrs. For cryo protection, a single plate-like crystal was transferred briefly to a drop containing 20% glycerol, 15% ethanol, 3% PEG 1K and 50 mM citrate/phosphate buffer pH 4.4, mounted in a loop, and flash-frozen in a 100 K nitrogen stream. A single dataset of 690° continuous phi measurement with 1° frames was collected at a wavelength of 1.54 Å on a Rigaku MicroMax-007HF X-ray generator equipped with VariMax optics Cu-HF and R-Axis IV++ image plate detector. The CysD1 structure was solved by sulfur single-wavelength anomalous diffraction (SAD) using the SHELX program (Sheldrick, 2015). The initial auto-built model was then iteratively rebuilt and refined using Coot and Phenix. A composite simulated annealing omit map was then produced and inspected to ensure the accuracy of the final model. Model geometry was assessed using Molprobity, and no Ramachandran outliers were detected.

#### VWF tubule modeling

A homology model of the VWF bead structure was generated by a combination of least squares fitting of known VWF structure fragments onto the MUC2 D1D2D3CysD1 bead and use of the Modeller software (Webb and Sali, 2016). The crystallographic structure of a monomeric mutant of the VWF D3 region (PDB code 6n29) (Dong et al., 2019) was divided into three segments spanning residues 764 to 827, 828 to 862, and 864 to 1241. Each segment was positioned by least squares fitting using Coot onto the appropriate region of the MUC2 bead. A homology model was generated using Modeller for the D1-D2 segment of VWF (residues 33 to 692). From this composite model the symmetry-related portion of the VWF bead was generated.

Modeling of the VWF tubule was done based on the published low resolution reconstruction of the VWF helical tubule (Huang et al., 2008). The VWF bead model was positioned with the two-fold axis of the bead perpendicular to the tubule axis and at a distance corresponding to the reported inner and outer diameters of the tubule (Huang et al., 2008). Helical symmetry was applied with 85.6° rotation and 26.2 Å rise, using Chimera (Pettersen et al., 2004). Rotations around the two-fold symmetry axis of the bead were tested in 2° steps to achieve the closest match with domain contours on the inner and outer faces of the low-resolution reconstruction of the VWF tubule (Huang et al., 2008), as the original maps were not available from the authors or from public repositories.

---

**Figure S1.**
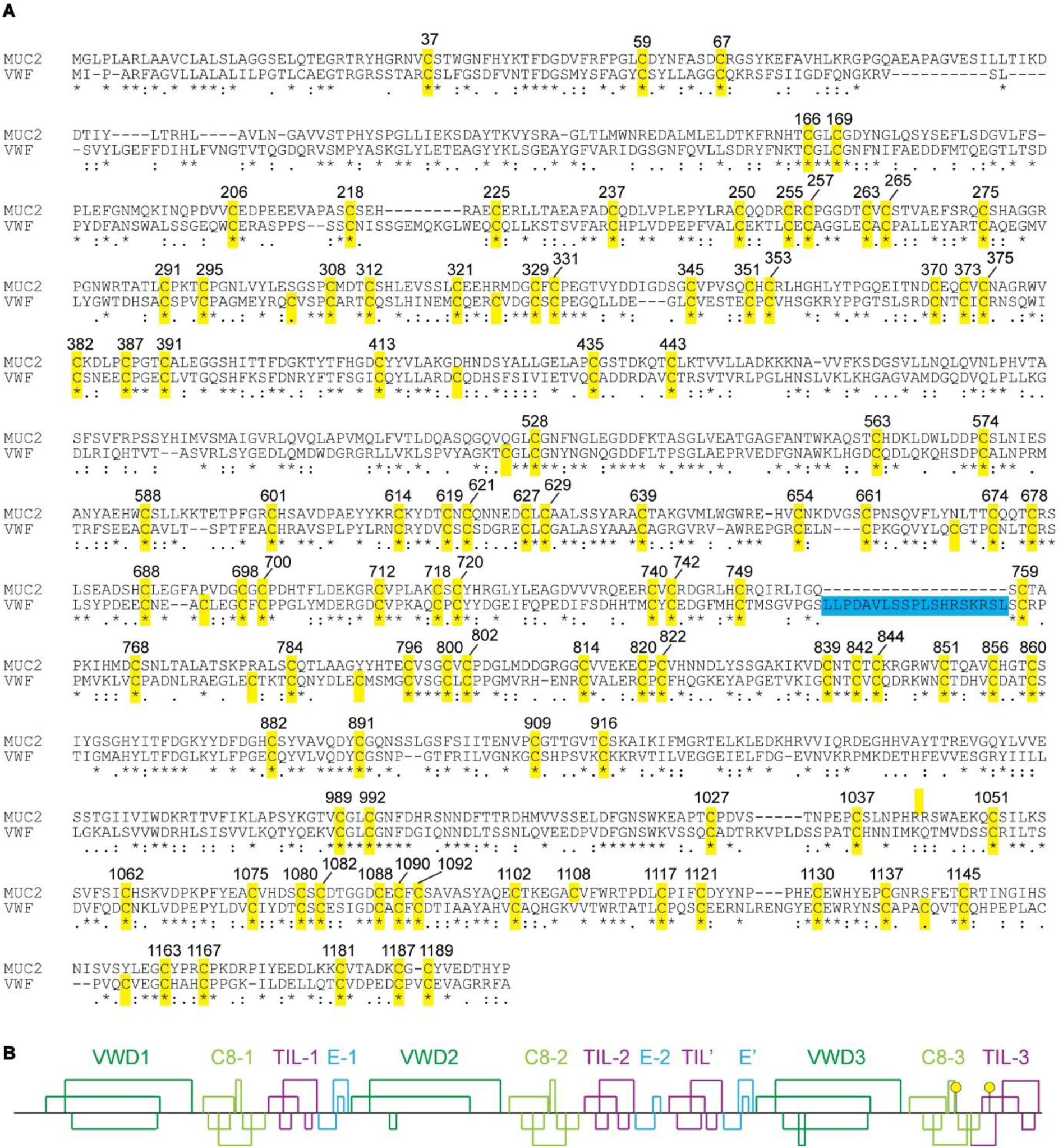
MUC2 disulfides and amino acid sequence alignment with VWF, related to Figure 1. (A) T-coffee (Notredame et al., 2000) was used to align the amino-terminal regions of human Mucin 2 (MUC2) and human von Willebrand factor (VWF). Cysteines (except in the signal peptides) are highlighted in yellow and labeled with numbers corresponding to the full-length MUC2 sequence (Uniprot Q02817). Highlighted in blue is an insertion in VWF that contains the furin cleavage site. (B) MUC2 disulfide bonding pattern mapped to scale. The first 1197 amino acids of MUC2 are shown. Cysteines that participate in intermolecular D3-D3 disulfide bonds are shown as yellow balls.

**Figure S2.**
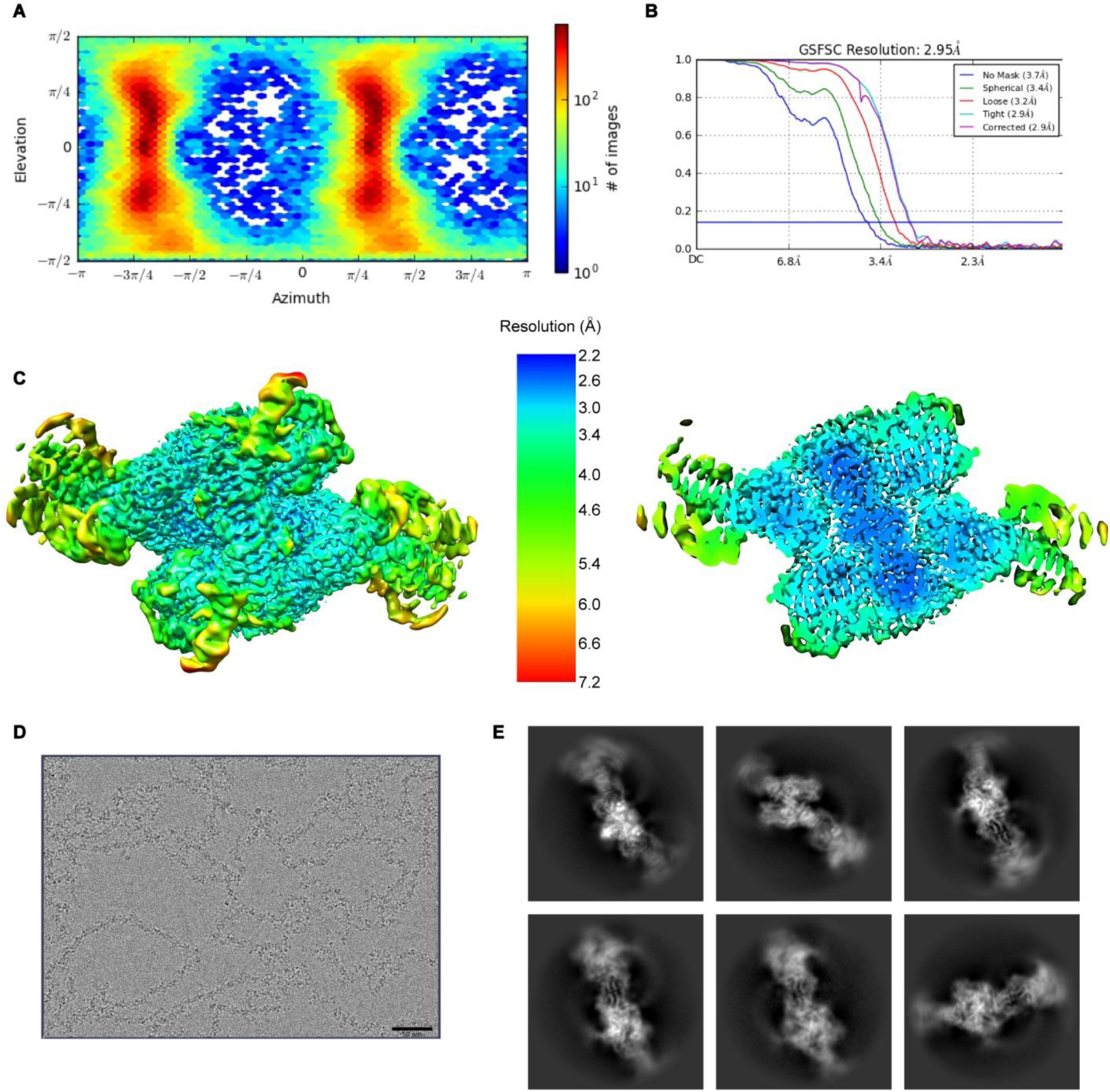
Single particle cryo-EM analysis of the MUC2 amino-terminal region polymer, related to Figure 3. (A) Angular distribution of particle projections. (B) Fourier shell correlation (FSC) curves. At a FSC 0.143 cut-off, the overall resolution for the map is 2.95 Å. (C) Map colored and filtered according to local resolution estimate, viewed along the 2-fold symmetry axis. A cutaway view is shown on the right. (D) Representative dose- and motion-corrected micrograph. Scale bar is 50 nm. Despite the frequent kinks in the chains, a predominant mode of interaction between adjacent beads was observed in the reconstructions. (E) Representative 2D classes showing the bead and the interface between beads from different angles and positions.

**Figure S3.**
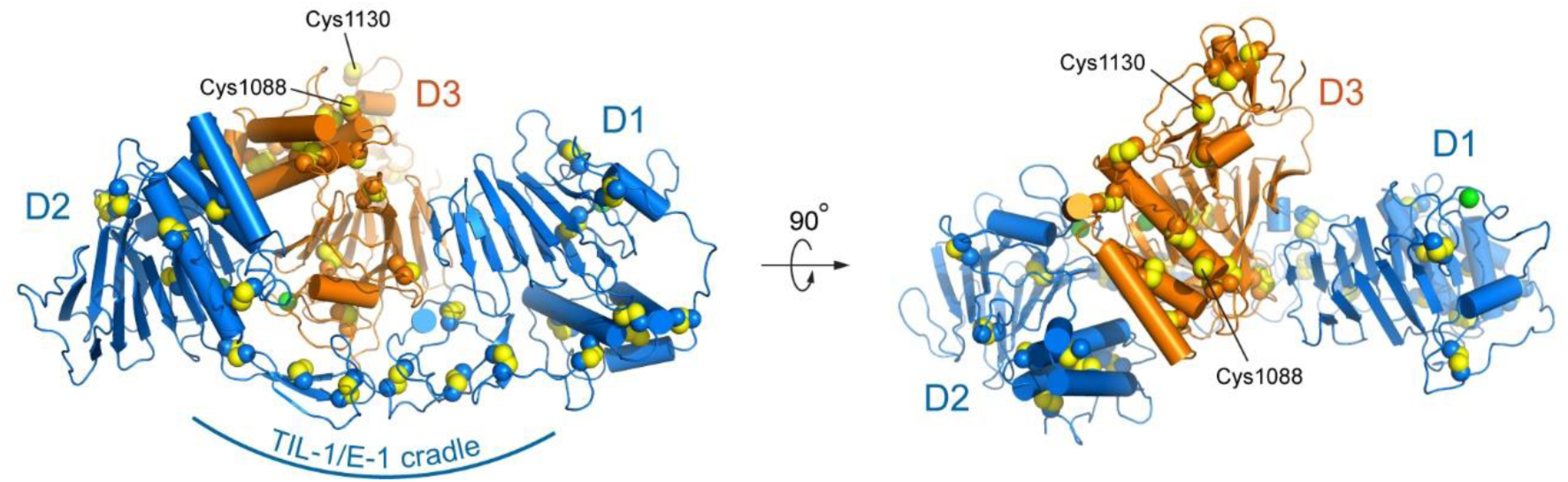
Orientation of D3 within the D1-D2 cradle, related to Figure 3. Coloring corresponds to Figure 3B and 3E. The cysteines (Cys1088 and Cys1130) that make intersubunit disulfide bonds are labeled. Based on the structure, these cysteines would be surface-exposed within the non-covalent dimer. However, it should be noted that when the VWF residue homologous to MUC2 1130 (VWF Cys1142) was mutated to alanine to form a monomeric version of the crystallized VWF D3 region (Dong et al., 2019), the surrounding polypeptide backbone was observed in a slightly different conformation than seen in this and previous (Javitt et al., 2019) disulfide bonded MUC2 structures. The VWF mutant configuration places residue 1142 near another disulfide bond in the structure. It is conceivable that the local geometry, or even the disulfide connectivity, changes along the assembly pathway through the ER and Golgi.

**Table S1.** Cryo-EM data collection, refinement and validation statistics, related to Figure 3.

|  |  |
| --- | --- |
|  | MUC2 D1D2D3CysD1<br>(EMDB-10517)<br>(PDB 6TM2) |
| **Data collection and processing** |  |
| Magnification | 105,000x |
| Voltage (kV) | 300 |
| Electron exposure (e-/Å<sup>2</sup>) | 40.6 |
| Defocus range (µm) | -1 to -2 |
| Pixel size (Å) | 0.85 |
| Symmetry imposed | C2 |
| Initial particle images (no.) | 1,430,962 |
| Final particle images (no.) | 178,136 |
| Map resolution (Å) | 2.95 |
| FSC threshold | 0.143 |
| Map resolution range (Å) | 2.2 to 7.2 |
| **Refinement** |  |
| Initial model used (PDB code) | *Ab initio*, PDB IDs 6RBF and 6TM6 |
| Model resolution (Å) | 2.7 (6RBF) and 1.6 (6TM6) |
| Map sharpening *B* factor (Å<sup>2</sup>) | Auto-estimated in Phenix |
| Model composition |  |
| Non-hydrogen atoms | 18,530 |
| Protein residues | 2,41 |
| Waters | 40 |
| Ligands | 30 |
| *B* factors (Å<sup>2</sup>) |  |
| Protein | 40.0 |
| Waters | 18.9 |
| Ligands | 75.4 |
| R.m.s. deviations |  |
| Bond lengths (Å) | 0.009 |
| Bond angles (°) | 1.38 |
| Validation |  |
| MolProbity score | 1.71 |
| Clashscore | 3.27 |
| Poor rotamers (%) | 0 |
| Ramachandran plot |  |
| Favored (%) | 88.4 |
| Allowed (%) | 11.2 |
| Disallowed (%) | 0.4 |

**Table S2.**
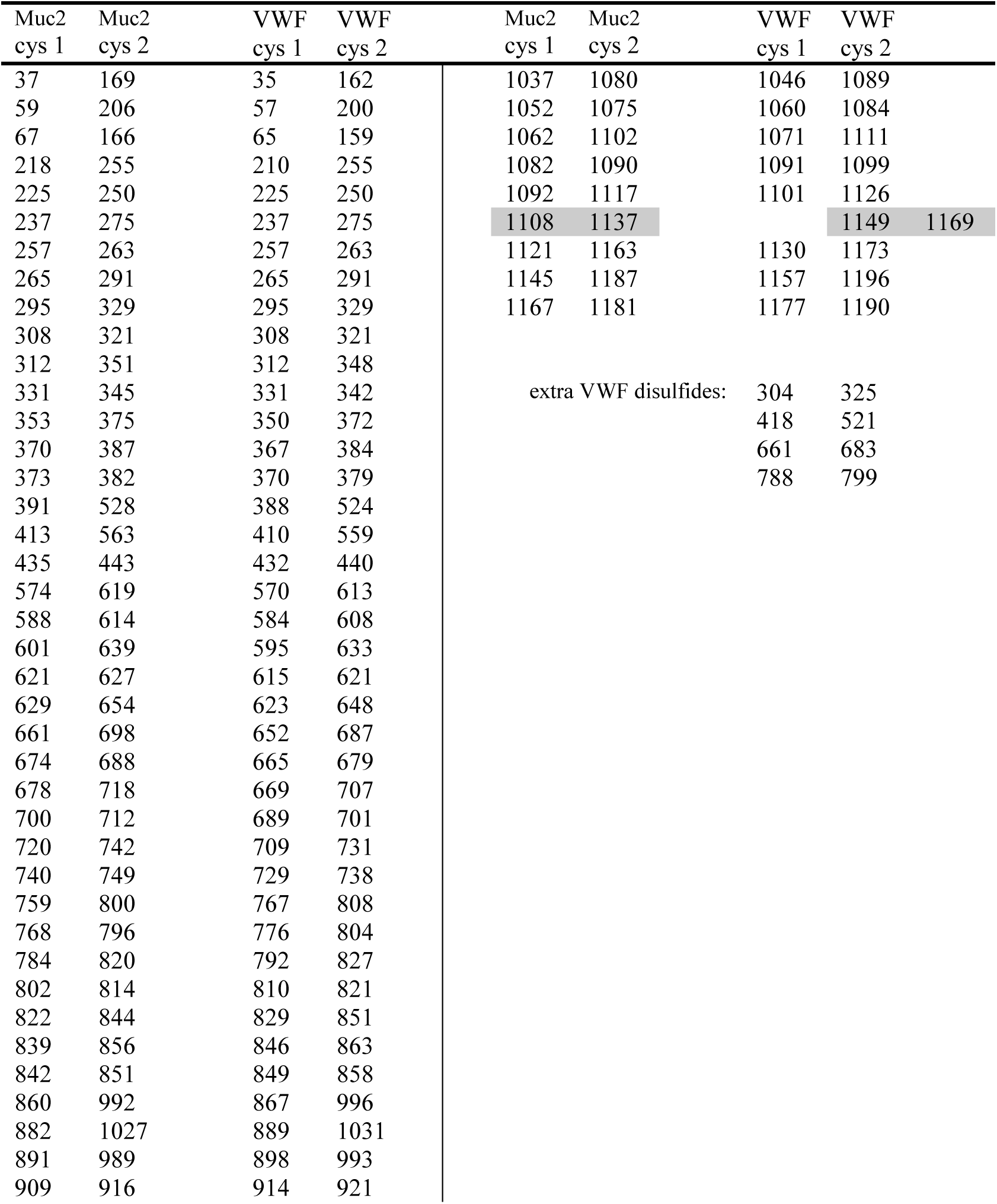
Disulfide connectivity of MUC2 and VWF, related to Figure 3. Numbers are according to the full-length sequences in UniProt (MUC2: Q02817; VWF: P04275). Each row shows disulfides that are homologous between MUC2 and VWF. Highlighted in gray is a pair of disulfides that share one conserved cysteine but a second divergent cysteine. Additional disulfides predicted in VWF due to close apposition of the corresponding residues in the homology model are listed as “extra VWF disulfides.” The VWF disulfide assignment is consistent with previous reports of Cys767-Cys808, Cys776-Cys804, and Cys810-Cys821 (Marti et al., 1987) and the TIL’E’ disulfides (Shiltagh et al., 2014).

**Table S3.**
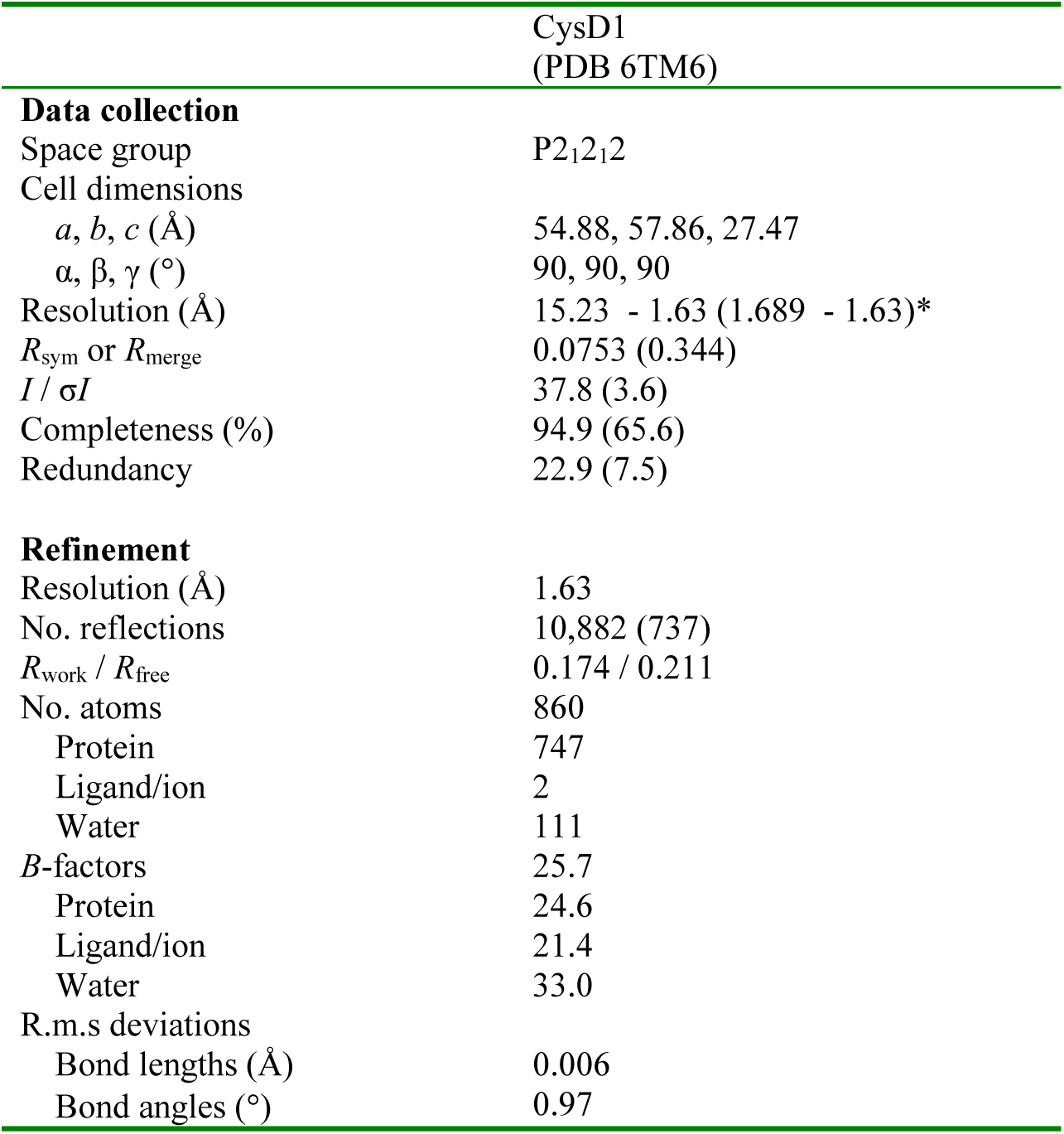
Crystallographic data collection, phasing and refinement statistics, related to Figure 5. Asterisk indicates the highest resolution shell, and data in parentheses correspond to this shell.

|  | CysD1<br>(PDB 6TM6) |
| --- | --- |
| **Data collection** |  |
| Space group | P2<sub>1</sub>2<sub>1</sub>2 |
| Cell dimensions |  |
| *a*, *b*, *c* (Å) | 54.88, 57.86, 27.47 |
| $\alpha$, $\beta$, $\gamma$ (°) | 90, 90, 90 |
| Resolution (Å) | 15.23 - 1.63 (1.689 - 1.63)* |
| *R*<sub>sym</sub> or *R*<sub>merge</sub> | 0.0753 (0.344) |
| *I* / σ*I* | 37.8 (3.6) |
| Completeness (%) | 94.9 (65.6) |
| Redundancy | 22.9 (7.5) |
| **Refinement** |  |
| Resolution (Å) | 1.63 |
| No. reflections | 10,882 (737) |
| *R*<sub>work</sub> / *R*<sub>free</sub> | 0.174 / 0.211 |
| No. atoms | 860 |
| Protein | 747 |
| Ligand/ion | 2 |
| Water | 111 |
| *B*-factors | 25.7 |
| Protein | 24.6 |
| Ligand/ion | 21.4 |
| Water | 33.0 |
| R.m.s deviations |  |
| Bond lengths (Å) | 0.006 |
| Bond angles (°) | 0.97 |

**Movie S1, related to Figure 3.**

Three beads of the MUC2 D1D2D3CysD1 polymer are displayed rotating around the polymer axis.

**Movie S2, related to Figure 5.**

Unfiltered cryo-EM map of MUC2 D1D2D3CysD1 demonstrates the arcs of density connecting the central bead with the two adjacent beads. The D1/D2 domains are colored orange and magenta, and the D3 segments are colored blue and green as in Figures 3E and 6C. CysD1 domains are colored cyan and light green. Adjacent beads and connecting arcs are gray. It is clear that CysD1 domains are connected to adjacent beads rather than to the D3 domains in the same bead. Correspondingly, the D3 domains within the central bead are linked to CysD1 domains in adjacent beads.

